# Establishment of a screening platform based on human coronavirus OC43 for the identification of microbial natural products with antiviral activity

**DOI:** 10.1101/2023.04.20.537680

**Authors:** Blanca Martínez-Arribas, Frederick Annang, Rosario Díaz-González, Guiomar Pérez-Moreno, Jesús Martín, Thomas A. Mackenzie, Francisco Castillo, Fernando Reyes, Olga Genilloud, Luis Miguel Ruiz-Pérez, Francisca Vicente, María C. Ramos, Dolores González-Pacanowska

## Abstract

Human coronaviruses (HCoVs) cause respiratory tract infections and are of great importance due to the recent severe acute respiratory syndrome coronavirus 2 (SARS-CoV-2) pandemic. Human betacoronavirus OC43 (HCoV-OC43) is an adequate surrogate for SARS-CoV-2 because it infects the human respiratory system, presents a comparable biology, and is transmitted in a similar way. Its use is advantageous since it only requires biosafety level (BSL)-2 infrastructure which minimizes costs and biosafety associated limitations. In this report, we describe a high-throughput screening (HTS) platform to identify compounds that inhibit the propagation of HCoV-OC43. Optimization of assays based on inhibition of the cytopathic effect and virus immunodetection with a specific antibody, has provided a robust methodology for the screening of a selection of microbial natural product extracts from the Fundación MEDINA collection. Using this approach, a subset of 1280 extracts has been explored. Of these, upon hit confirmation and early LC-MS dereplication, 10 extracts were identified that contain potential new compounds. In addition, we report on the novel antiviral activity of some previously described natural products whose presence in bioactive extracts was confirmed by LC/MS analysis.

**IMPORTANCE:** The COVID-19 pandemic has revealed the lack of effective treatments against betacoronaviruses and the urgent need for new broad-spectrum antivirals. Natural products are a valuable source of bioactive compounds with pharmaceutical potential that may lead to the discovery of new antiviral agents. Specifically, compared to conventional synthetic molecules, microbial natural extracts possess a unique and vast chemical diversity and are amenable to large-scale production. The implementation of a high-throughput screening platform using the betacoronavirus OC43 in a human cell line infection model has provided proof of concept of the approach and has allowed for the rapid and efficient evaluation of 1280 microbial extracts. The identification of several active compounds validates the potential of the platform for the search for new compounds with antiviral capacity.

## INTRODUCTION

Coronaviruses (CoVs) have the ability to propagate and generate new species that cause epidemic diseases. The emergence of CoV infections has persistently caused serious public health alarms over the years. Severe acute CoV infections, including the respiratory syndrome-related coronavirus (SARS-CoV), the Middle East respiratory syndrome-related coronavirus (MERS-CoV), and the pandemic virus SARS-CoV-2, have become a rising and long-lasting global risk. The impact of the SARS-CoV-2 pandemic has revealed the urgent need for broad-spectrum antivirals to prevent future viral pandemics of unknown origin.

To date, efficient vaccines are lacking for many viruses and novel antiviral drugs with clinical efficacy are called for. The situation is further aggravated by the potential development of mutants that are drug resistant, thus limiting drug efficacy (1-4)

Natural products represent a rich source of diverse, new, and inexpensive chemical starting points for drug discovery in the area of antivirals and serve as an outstanding source of biodiversity for the discovery of novel compounds. Indeed, approximately 65% of successfully approved drugs are either natural products or derivatives (5). Many natural products derived from microbes have been reported to have robust antiviral activity, with anti-hepatitis, anti-herpes simplex, anti-HIV, anti-influenza, anti-respiratory syncytial virus, and anti-SARS-CoV-2 properties (6-12). However, when performing the search of active compounds from whole crude microbial extracts, the process is both time consuming and capital intensive. For that reason, it is very critical to develop quick, efficient, and high-quality high-throughput primary screening assays integrated with chemistry platforms for early dereplication of known compounds to guarantee that only extracts with relevant pharmacological properties are selected for further development downstream.

Antiviral drug discovery using SARS-CoV-2 for quantitative screening requires a biosafety level (BSL)-3 facility, which is costly and labour-intensive. Hence, the availability of model systems of HCoV for high-throughput screening (HTS) assays that involve BSL-2 is highly advantageous. Human coronavirus OC43 (HCoV-OC43) (order: Nidovirales, family: Coronaviridae, genus: Betacoronavirus, subgenus: Embecovirus, species: Betacoronavirus1) exhibits several common features with SARS-CoV-2 regarding structure and biology, which makes it a well-matched surrogate for SARS-CoV-2 (13). As members of the same genus, HCoV-OC43 and SARS-CoV-2 are closely related genetically (14) and they are transmitted by aerosols and droplets and replicate in human respiratory epithelium (15).

A wide variety of methods to evaluate HCoV-OC43 infection using different cell lines have been described (13). With regards to host cells, HRT-18 (human colon cancer cells) (16, 17), MRC-5 (human lung fibroblasts (18), Vero E6 expressing TMPRSS2 (19) among others have been utilized. In addition, different detection methods (plaque assay, indirect immunostaining, resazurin reduction) (13, 20) and infection conditions have been explored. Here we report the standardisation of methodology for the quantification of infection by HCoV-OC43 and describe the development of a robust experimental approach using a HTS platform to identify compounds that exhibit anti-OC43 activity. The experimental design has been successfully applied to analyse 1280 microbial extracts from actinomycetes and fungi of the Fundación MEDINA’s library (Supplemental Figure 1). We have combined the use of determination of the cytopathic effect (CPE) with antibody-based detection bioimaging assays in the analysis which revealed the presence of novel antiviral compounds. To our knowledge this is the first report of HTS using microbial natural product extracts against HCoV-OC43.

## MATERIALS AND METHODS

### Reagents and antibodies

Dimethyl sulfoxide (DMSO), chloroquine diphosphate salt, mycophenolate mofetil, linoleic acid, destruxin A, bovine serum albumin (BSA) and resazurin were purchased from Sigma Aldrich. Tamoxifen, ribavirin, destruxin B and gymnoascolide A were purchased from Tocris, Santa Cruz Biotechnology, Adooq and Cayman, respectively. All compounds were dissolved in 100% DMSO. The mouse monoclonal anti-HCoV-OC43 antibody (MAB2012) was purchased from Millipore, the secondary antibody AlexaFluor® 488-conjugated anti-mouse and Hoechst 33342 were purchased from Thermo-Scientific.

### Cell culture

The human lung fibroblasts cell line MRC-5 (CCL-171, ATCC) and the cell line BHK-21 (CCL-10, ATCC) were cultured in Minimum Essential Medium (MEM) (Life Technologies) supplemented with 10% fetal bovine serum (FBS) (Life Technologies) and 100 units/mL penicillin and 100 μg/mL streptomycin (Life Technologies). Cells were incubated at 37 °C in a humidified atmosphere of 5% CO_2_ and were periodically analysed and confirmed to be mycoplasma negative.

### Virus production

The human betacoronavirus HCoV-OC43 (VR-1588, ATCC) was propagated in MRC-5 cells. MRC-5 cells at a 90% confluence were inoculated with HCoV-OC43 in infection media (MEM, 2% inactivated FBS, Pen/Strep) and incubated 2 h at 33 °C, rocking the flask every 15 min for virus adsorption. After virus adsorption, infection media was added, and infected cells were incubated at 33 °C for 5 to 7 days until more than 50% of the cells presented a cytopathic effect (CPE) resulting in cell death. For virus recovery, an infected culture was subjected to 3 freeze-thaw cycles, centrifuged at 500 g, 10 min at 4 °C to spin down cells and cell debris. Viral particles were recovered from supernatant and aliquoted in cryotubes. Viral stock aliquots were then rapidly frozen in a dry-ice/ethanol bath and stored at −80 °C.

### Batch infection with HCoV-OC43

MRC-5 and BHK-21 cells were seeded 24 h prior to infection for a 90% confluence at the time of infection. Different multiplicities of infection (MOI) were tested for HCoV-OC43 infection of MRC-5 and BHK-21 cells. The virus adsorption was performed for 2 h at 33 °C, rocking the cells every 15 min, and then infected cells were incubated 24 h at 33 °C before seeding into 96 or 384 well plates.

### Resazurin-based assay

Assay plates were previously prepared with microbial extracts to be screened at a final concentration of 1/100 dilution of whole broth equivalent (WBE) and at a final 0.2% DMSO concentration in 96 well plates. Infected cells were washed, trypsinized and seeded in assay plates at a cellular concentration of 2×10^4^ cells/well in infection media. The plates were incubated at 37 °C for 96 h in the presence of the extracts. 5 days after infection media was withdrawn and 120 μL of infection media containing 20% resazurin (Sigma-Aldrich) was added per well. Plates were incubated for 2 h at 37 °C and fluorescence was determined at 550-590 nm in a TECAN infinite F200 plate reader. Non-infected cells and infected cells with 0.2% DMSO were used as positive and negative control, respectively. Extracts showing a CPE inhibition superior to 40% were selected as hits.

The resazurin assay was also used to determine the cytotoxicity of extracts or compounds in MRC-5 cells. Briefly, 2×10^4^ cells /well were seeded in 96 well plates containing the microbial extracts or pure compounds. After 96 h cells culture media was withdrawn and 120 μL of infection media containing 20% resazurin (Sigma-Aldrich) was added per well. Plates were incubated for 2 h at 37 °C and fluorescence was determined at 550-590 nm in a TECAN infinite F200 plate reader. MRC-5 cells treated with 50 μM of tamoxifen were used as negative control, while positive controls corresponded to MRC-5 cells incubated in presence of 0.2% DMSO for 96 h.

### High content screening assay

Assay plates were previously prepared with microbial extracts to be screened at a final concentration of 1/100 dilution of whole broth equivalent (WBE) and at a final DMSO concentration of 0.2% in 384 well black bottom-transparent plates (Greiner bio-one). MRC-5 infected cells were washed, trypsinized and seeded in assay plates at a cellular concentration of 3000 cells/well in infection media. The plates were incubated at 37 °C for 72 h in the presence of the extracts. 4 days post-infection cells were subjected to indirect immunostaining by using an anti-OC43 monoclonal antibody (MAB2012). Briefly, cells were washed with 1× PBS, fixed with 4% *p-*formaldehyde and permeabilized with 0.4% Triton X-100. Next, cells were incubated in 5% BSA blocking solution for 2 h and then incubated with the anti-HCoV-OC43 monoclonal antibody at a 1:2000 dilution O/N at 4 °C. Cells were washed, incubated with antibody AlexaFluor® 488-conjugated anti-mouse, washed again and nuclei were stained with 4 μM Hoechst 33342. Digital images were captured using Operetta CLS High Content Analysis System (PerkinElmer) with a 5× air objective. Images were analysed using the Harmony software (PerkinElmer) and the following parameters were determined: number of nuclei per well, number of infected cells per well and total 488-fluorescence signal per well. Based on the 488 signal and cell nuclei, cells are segmented for the calculation of the total 488-fluorescence signal.

### LC-MS dereplication and database matching of known secondary metabolites

The active extracts of interest were subjected to LC-MS dereplication and database matching to identify possible known components using an Agilent (Santa Clara, CA) 1260 Infinity II single Quadrupole LC-MS system and LC/MS analytical conditions and methodology previously described (21-23)

### RNA isolation and Real-time RT-PCR

Viral RNA from supernatants was purified using the Macherey-Nagel Nucleospin RNA Kit. RT-qPCR was performed in a single-step using the One-Step TB Green PrimeScript RT-PCR Kit II (Takara Bio). The HCoV-OC43 nucleocapsid gene was amplified with the following primers: forward primer, 5’ AGCAACCAGGCTGATGTCAATACC-3’ and the reverse primer, 5’ AGCAGACCTTCCTGAGCCTTCAAT-3’(24). A standard curve was generated with purified HCoV-RNA (Vircell).

### Data analysis

#### CPE inhibition

Extract activities were calculated automatically using the Genedata Screener software (Genedata AG, Basel, Switzerland), and the percentage of CPE inhibition of each extract was determined by equation 1 integrated in the Genedata Screener software:

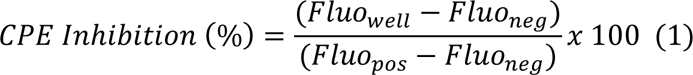

where Fluo_well_ is the measured fluorescence for each well; Fluo_pos_ is the average fluorescence for the positive control (infected MRC-5 cells treated with 400 μM ribavirin) and Fluo_neg_ is the average fluorescence for the negative control (infected MRC-5 cells 0.2% DMSO).

#### Cytotoxicity

Cellular cytotoxicity was determined by equation 2:

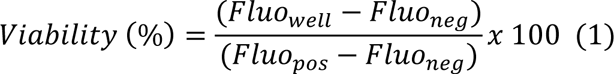

where Fluo_well_ is the measured fluorescence for each well; Fluo_neg_ is the average fluorescence for the negative control (MRC-5 cells treated with 50 µM tamoxifen) and Fluo_pos_ is the average fluorescence for the positive control (MRC-5 cells 0.2% DMSO).

#### HCoV-OC43 infection

The sum intensity (SumInt) per well corresponding to the Alexa 488 fluorescence signal was automatically given by the Operetta Analysis System software. HCoV-OC43 infection percentages were determined by equation 3:

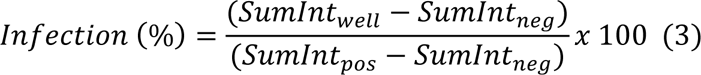

SumInt_well_ corresponds to the sum of 488-fluorescence signal in wells with extract and SumInt_neg_ and SumInt_pos_ correspond to the 488-fluorescence signal of negative and positive controls wells, respectively. Positive control corresponds to MRC-5 infected cells with 0.2% DMSO and the negative control refers to MRC-5 infected cells treated with 400 µM ribavirin.

The 50% cytotoxic concentrations (CC_50_) and the half-maximal effective concentration (EC_50_) were determined using SigmaPlot 15.0 software.

#### Evaluation of quality control (QC) of the screening assay

Z’-Factor and signal to background (S/B) are quality control parameters that reflect the robustness and reproducibility of the assay. Z’-Factor is a statistic parameter used to evaluate the quality of an assay and it is calculated by relating the sum of the standard deviations for the reference wells to the signal range given by the difference in their mean values:

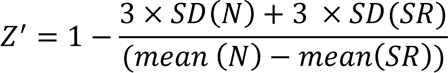

Being SD: standard deviation, N: neutral controls and SR: scale reference. A value of 0.4 - 1 characterizes a robust assay. S/B is the ratio of the mean value of the raw data for positive and negative controls.

## RESULTS

### Set up of a resazurin-based assay for the detection of antivirals that inhibit CPE

In order to optimize the protocol of batch infection with HCoV-OC43, different MOIs were tested in two different cell lines that had been previously described as host models for HCoV-OC43: MRC-5 and BHK-21 (17, 25). Both cell lines were infected with increasing HCoV-OC43 MOIs (0.01, 0.1 and 0.5) and 24h post-infection cells were seeded in 96 well plates. CPE-induced cell death was assessed 5 days post-infection using a resazurin-based approach. In this assay the fluorescence signal is directly proportional to cell survival and accordingly decreases when the viral CPE causes cell death. For both cells lines we obtained a reproducible and homogenous infection in the 96 well plate format at MOIs higher than 0.1. A decrease in viability of BHK-21 and MRC-5 cells was not observed at the lower MOI of 0.01 (Figure 1A).

**Figure 1:**
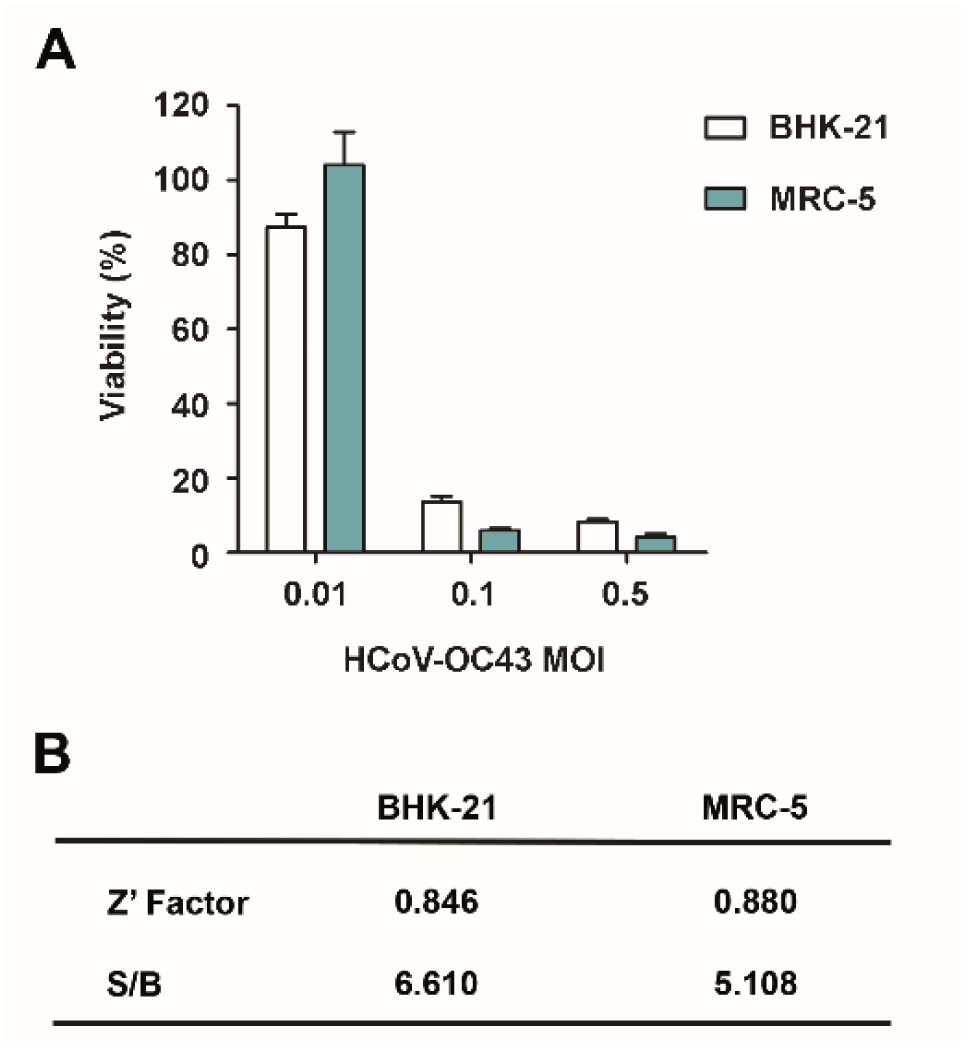
CPE evaluation. A. Cell viability of MRC-5 and BHK-21 cell lines 5 days post-infection with HCoV-OC43 at different MOIs, 0.01, 0.1 and 0.5. B. Average Z’factor and signal to background (S/B) values for BHK-21 and MRC-5 cells infected with HCoV-OC43.

MRC-5 cells infection at a 0.1 MOI was selected as the most suitable model, since the decrease in cell viability 5 days post-infection was higher than 90%, while in the case of BHK-21 cells with the same MOI, the loss of cell viability was less pronounced (80% decrease). Z’ factors were 0.88 for MRC-5 cells and 0.84 for BHK-21 emphasizing the robustness of the assay in both cases (Figure 1B).

For the selected model, the progression of the CPE up to 5 days post-infection gives rise to a cell death of MRC-5 infected cells higher than 90%. This CPE will be inhibited in the presence of active antiviral components and thus the resazurin-based fluorescence signal obtained will increase and be compared to the signal corresponding to non-infected cells (100% viability control). Pharmacological validation of the assay was carried out using three compounds that had been previously described as antivirals against SARS-CoV-2 and HCoV-OC43: ribavirin, chloroquine and mycophenolate mofetil (25-28).

The EC_50_ values were determined for the three compounds together with the corresponding CC_50_ values in non-infected MRC-5 control cells (Figure 2). Chloroquine showed a low EC_50_ of 6.22 μM, although it was also the most cytotoxic with a CC_50_ of 14.8 μM. Ribavirin and mycophenolate mofetil showed no cytotoxicity at concentrations below 800 μM and 200 μM, respectively. As for the antiviral activity, mycophenolate mofetil exhibited a lower EC_50_ (20.66 μM) compared to ribavirin (118.38 μM).

**Figure 2:**
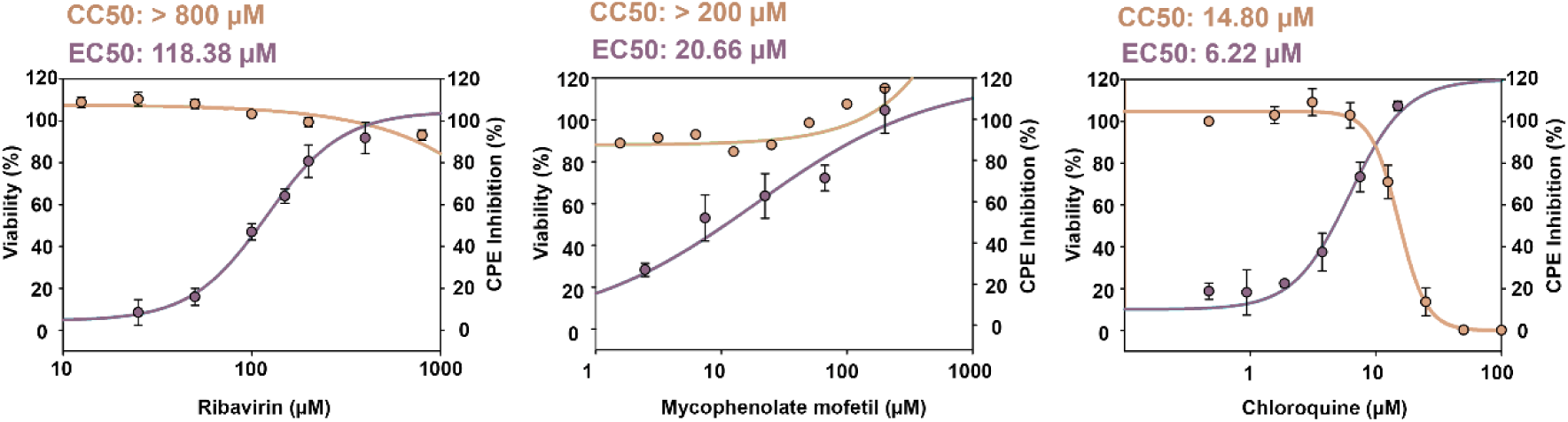
Evaluation of reference antivirals ribavirin, mycophenolate mofetil and chloroquine. Determination of EC_50_ and CC_50_ values after 96 h incubation with the drugs of infected cells (after 5 days of infection) and non-infected cells respectively. EC_50_ and CC_50_ values were calculated using SigmaPlot software.

Ribavirin was selected as the antiviral control compound to be routinely included in the assay. A concentration of 400 μM of ribavirin was used, since it showed a good selectivity index (SI), close to 8, and it has been widely described and used as broad-spectrum antiviral (29).

### Antiviral primary screening and hit identification

For the primary screening of 1280 (320 fungal, 960 actinomycete) microbial extracts from the MEDINA microbial library, the resazurin-based assay for CPE inhibition determination was used (general workflow is shown in Figure 3). The average Z’ factor for the assay was 0.971 ± 0.04, and the fluorescence signal of infected cells was 5 times lower than the positive control (Figure 4A and B). We selected as hits those extracts that inhibited more than 40% the CPE (Figure 4C). Out of the 1280 microbial extracts, 40 were selected as hits (a hit rate of 3.12%) (Table 1); 13 corresponded to actinomycete extracts and 27 corresponded to fungal extracts.

**Figure 3:**
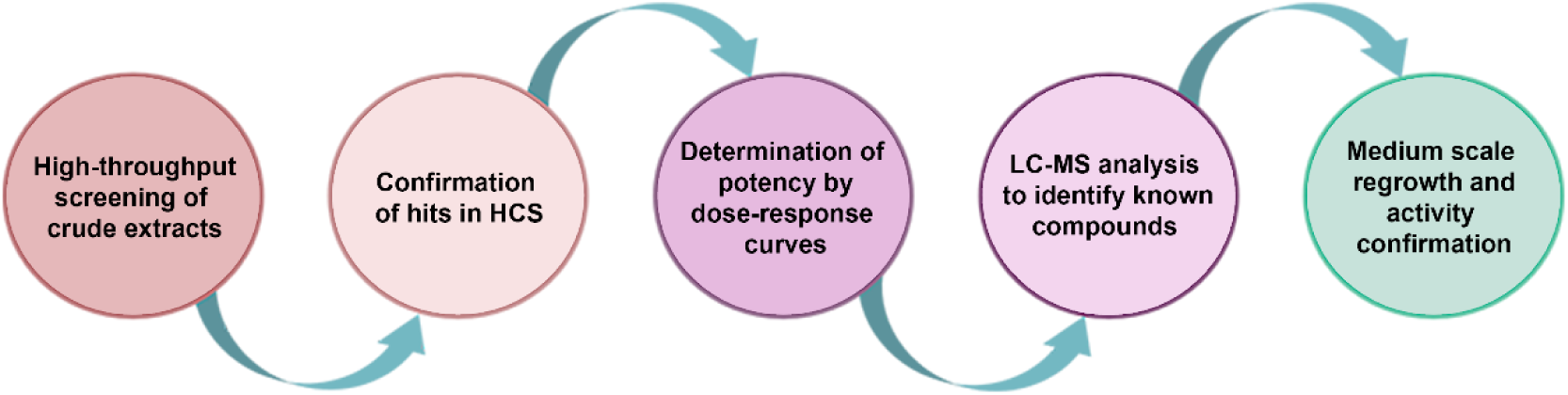
Schematic representation of the platform for the discovery of antiviral drugs from microbial extracts.

**Figure 4.**
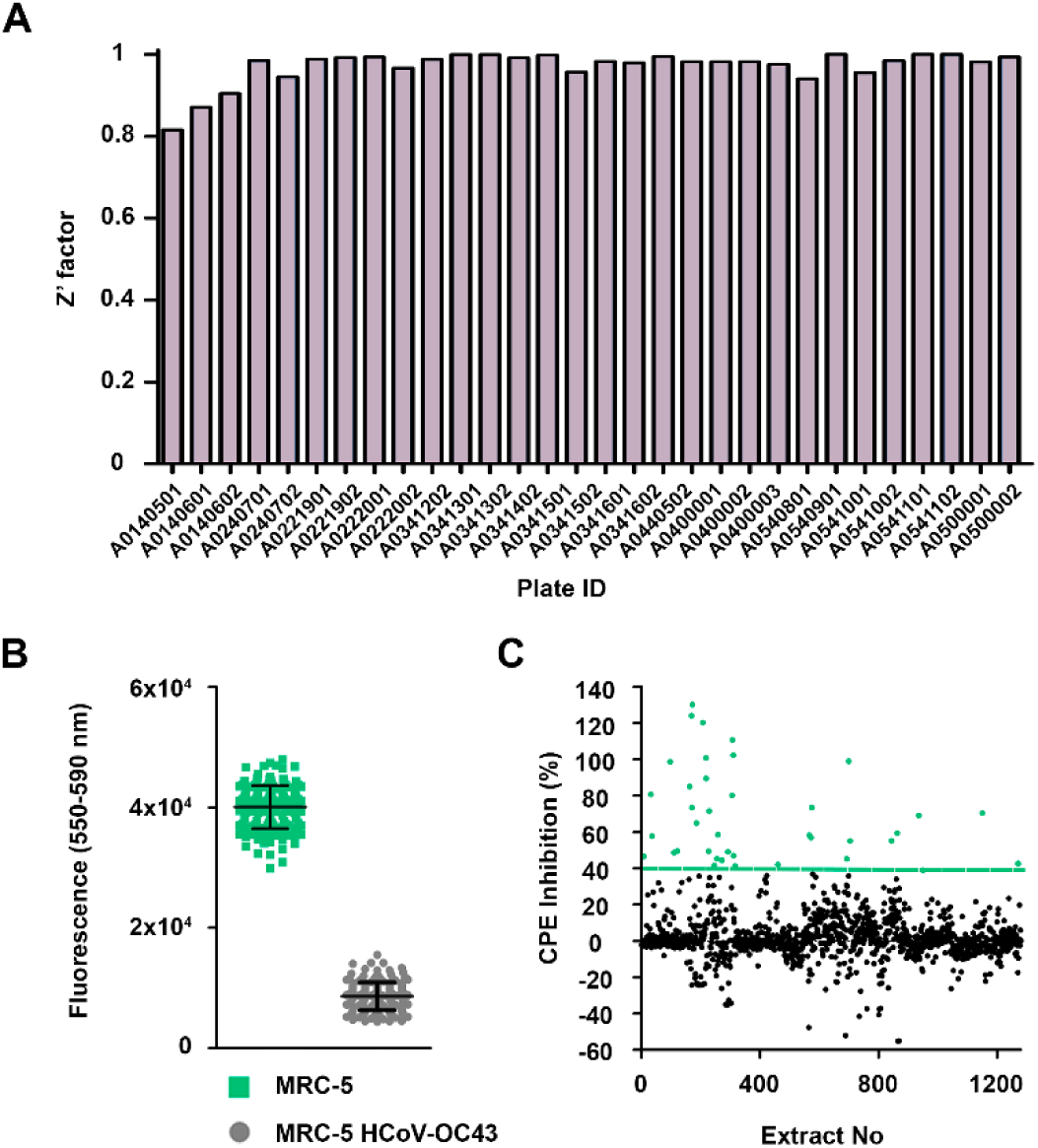
Method validation and primary screening of a subset of 1280 microbial extracts. A. Distribution of the Z’ factor along the different plates. B. Fluorescence distribution of the positive control (MRC-5 infected with 0.1 MOI HCoV-OC43 and treated with 400 μM ribavirin) and the negative control (MRC-5 infected with 0.1 MOI HCoV-OC43). C. Representation of 1280 extracts from primary screening. The green line represents the cut-off for hit selection (40% inhibition of CPE) and the circles in green correspond to the 40 selected hits presenting a CPE inhibition higher than 40%.

**Table 1.** Number of extracts identified as hits at different stages of the screening process.

| Screening stages | Number of active extracts (%) |
| --- | --- |
| Primary screening | 40 (3.12) |
| Cherry-picking for confirmation HCS | 32 (2.50) |
| Dose-response curves for EC<sub>50</sub> calculation | 20 (1.56) |
| Prioritized for regrowth in 100 mL volume | 15 (1.17) |
| Activity reconfirmation after 100 mL regrowth | 12 (0.93) |
| Dose-response curves for EC<sub>50</sub> calculation after 100 mL regrowth | 10 (0.78) |

### Implementation of a high-content screening in 384-well format

In the CPE inhibition screening, cytotoxicity can interfere with the assay since the readout is the same for CPE-induced death and toxicity caused by the extracts. High content screening (HCS) is an approach that combines automated imaging and quantitative data analysis in a high-throughput format and allows to simultaneously evaluate different parameters. We successfully optimized HCS methodology in a 384-well format for the parallel determination of antiviral activity and toxicity of microbial extracts by a dual labelling of HCoV-OC43 and cell nuclei (Figure 5A). The infection conditions were maintained between the two methodologies, with the exception of the experiment endpoint, that was reduced to 4 days post-infection for HCS since we are no longer quantifying CPE/death inhibition but viral presence in viable cells. To optimize the methodology, three different densities of MRC-5 batch-infected cells (MOI 0.1) were assayed in the 384-well format. A density of 4000 infected cells per well was selected. Z’ factor for this assay was 0.88 and a signal to background (S/B) window of 21.79 compared to the S/B of 5.11 from the CPE inhibition assay.

**Figure 5.**
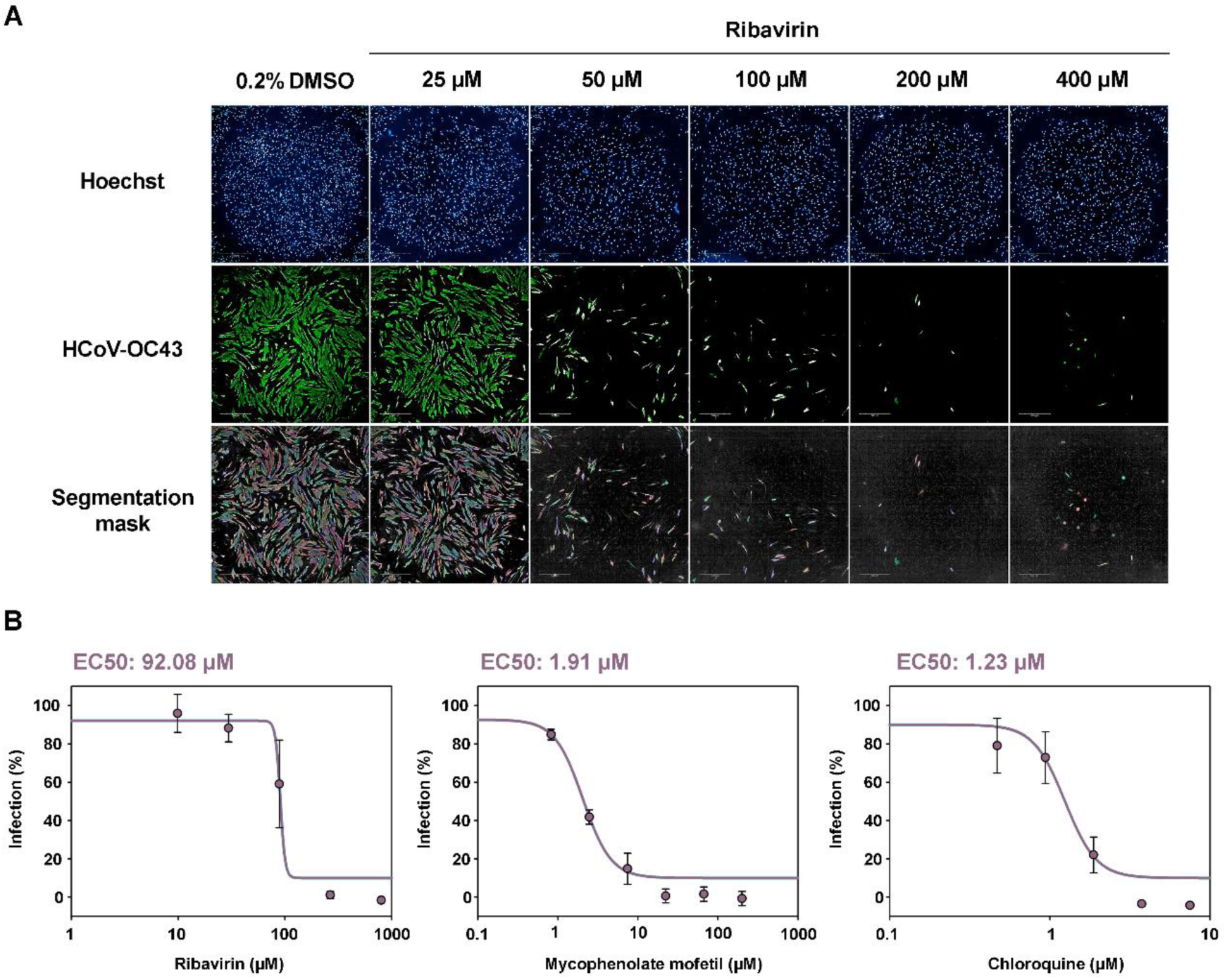
Set up of the high content screening assay with reference antivirals. A. Immunofluorescence of infected cells treated with increasing concentration of the reference compound ribavirin. B. Determination of EC_50_ values for ribavirin, mycophenolate mofetil and chloroquine after 72 h incubation with the drugs and 4 days of infection. EC_50_ values were calculated using SigmaPlot software.

We validated the HCS model with the three reference compounds previously assayed: ribavirin, chloroquine and mycophenolate mofetil. EC_50_ values were mostly consistent between both methodologies, except for mycophenolate mofetil that showed a much lower EC_50_, 1.21 μM, compared to 20.66 μM determined in the CPE inhibition assay (Figure 5B) exhibited again the lowest. The EC_50_ obtained this way for chloroquine was 1.23 μM and for ribavirin 92.08 μM.

### Hit confirmation

We assessed the activities of the extracts selected in the CPE inhibition assay and confirmed their antiviral activity by cherry-picking. The average Z’ factor was 0.780 ± 0.09 and the fluorescent signal in infected cells when compared with infected cells treated with 400 μM ribavirin was 30 times higher, improving the difference observed in the CPE assay (Figure 6A and B). We established a cut-off value of lower than 70% infection for hit confirmation (Figure 6C). Out of the 40 hits, 32 (9 actinomycetes and 23 fungi) confirmed their antiviral activity (Table 1) using the immunodetection bioimaging assay with a specific anti-HCoV-OC43 antibody. The high rate of hit confirmation using two independent assays further reinforces the robustness of the approach. Dose-response experiments were subsequently performed to evaluate extract potency. Dose-response curves consisted in 5 points of serial 2-fold dilutions were assayed in duplicate (Figure 7). Twenty extracts presented an EC_50_ value lower than 0.7 whole broth equivalents (WBEs) and were selected and progressed for liquid chromatography mass spectrometry (LC-MS) dereplication.

**Figure 6.**
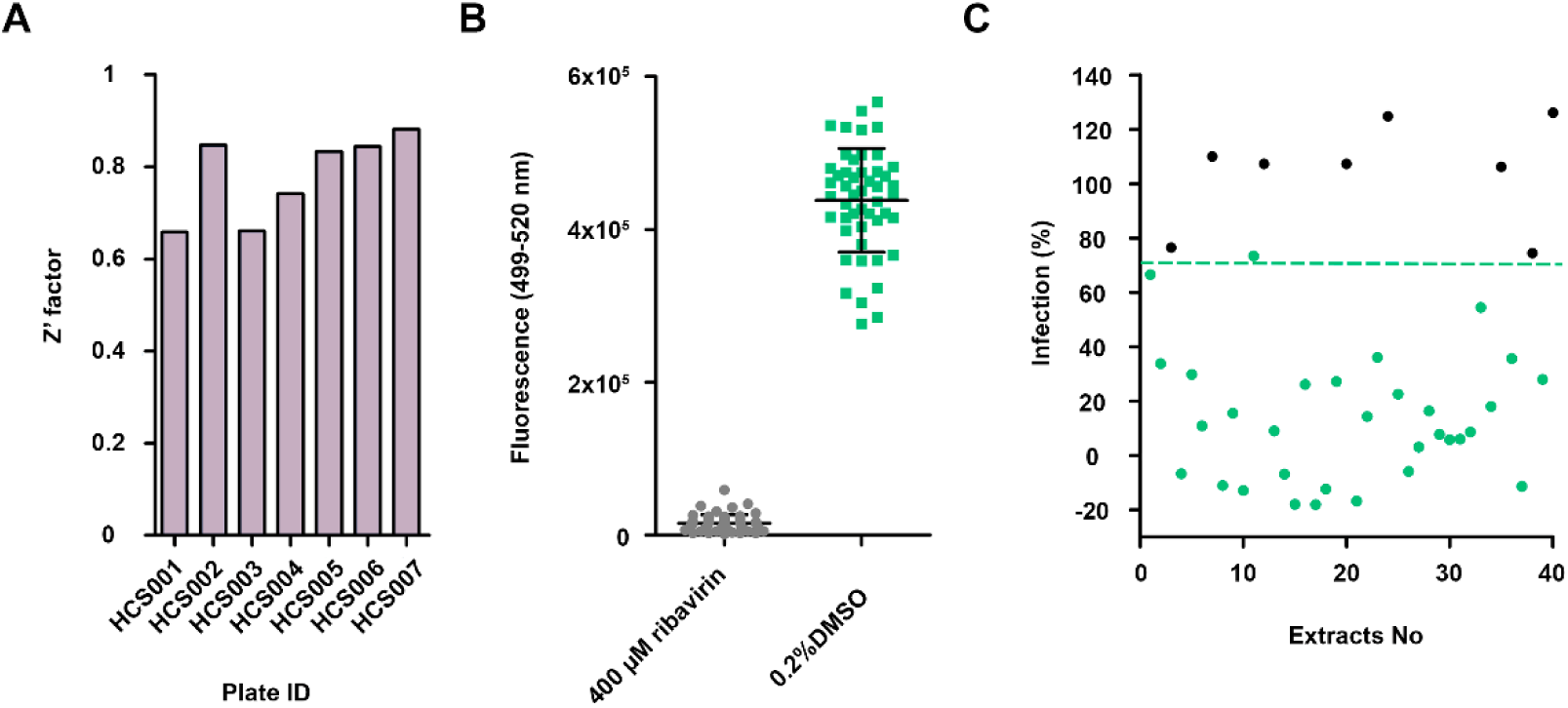
High content screening assay validation. A. Distribution of the Z’ factor along the different plates. B. Distribution of the fluorescence signal of the negative control (infected cells treated with 400 μM of ribavirin) and the positive control (infected cells with 0.2% DMSO). C. Representation of the 40 hits from the primary screening. The green line represents the cut-off for hit selection (70% of infection) and the circles in green correspond to the 32 extracts that presented a percentage of infection, compared to controls, lower than 70%.

**Figure 7.**
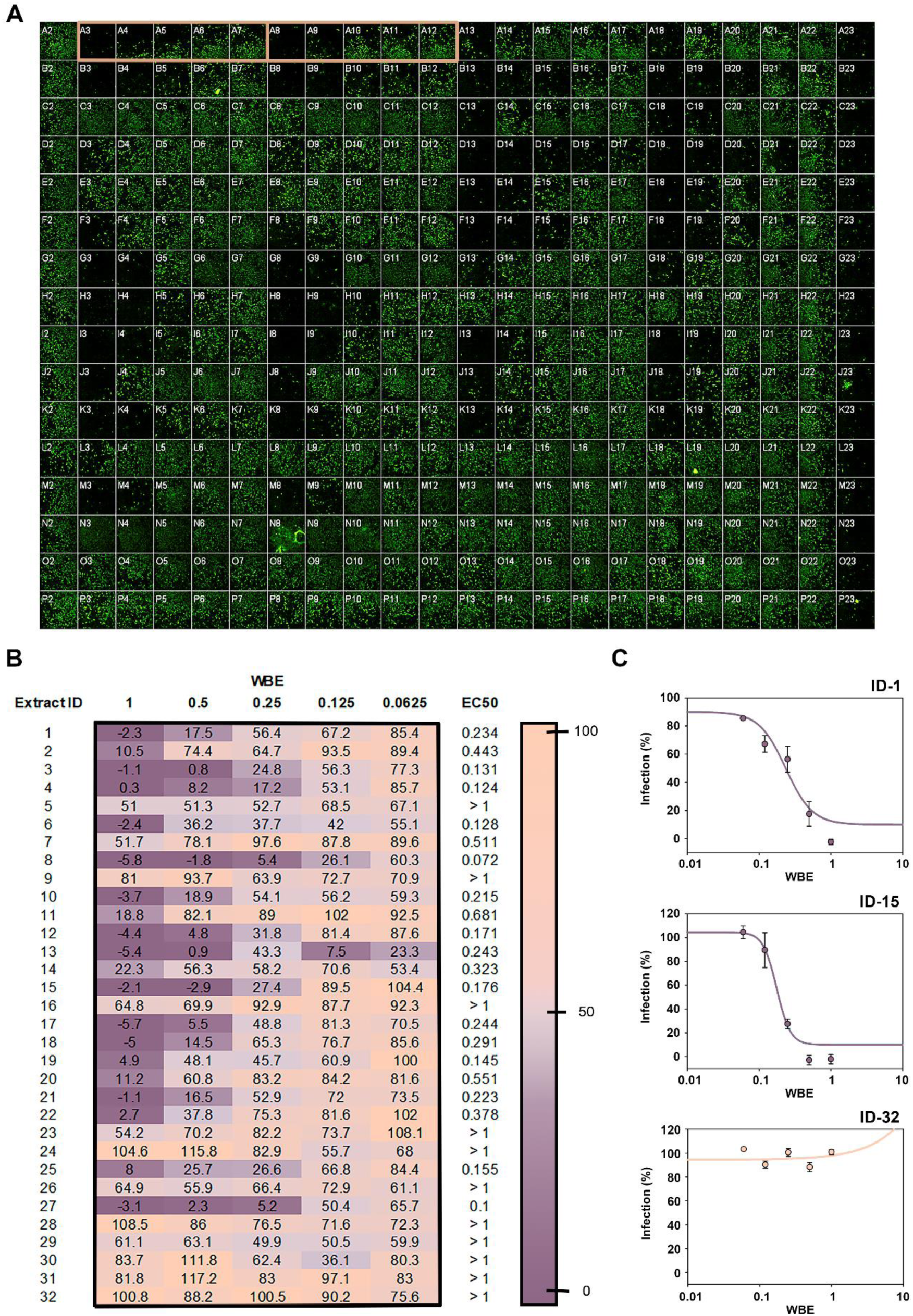
Dose-response curves for potency determination. A. Visualization of an HCS-384-well plate. Column 1 (from A2 to P2) corresponds to the 100% infection control: MRC-5 cells infected with HCoV-OC43 0.2% DMSO. Column 22 (from A23 to P23) corresponds to the 0% infection control: MRC-5 cells infected with HCoV-OC43 treated with 400 μM ribavirin. Columns 2 to 21 contain five points of 2-fold dose-response curves of extracts. Example of one of the dose-response curves in duplicate is squared in orange. B. Heatmap of the infection percentage in dose-response curves of the 32 hits. EC_50_ values are expressed as WBEs. C. Examples of dose-response curves of extracts. ID-1 and ID-15 correspond to active potent extracts (EC_50_ < 0.7 WBE) while ID-32 corresponds to a non-potent extract (EC_50_ > 0.7 WBE). EC_50_ values were calculated using SigmaPlot software.

### Early LC-MS dereplication and bioassay-guided fractionation

LC-MS analysis allowed for the detection and identification of the main components of active extracts. A MEDINA’s in-house application to search for matches of the UV-LC-MS data of the metabolites found in the active extracts to known metabolites already registered in our database (22) was used. It was identified 19 known compounds (Figure 8A) present in the bioactive extracts and purchased those commercially available: destruxin A, destruxin B, linoleic acid and gymnoascolide A. The four compounds were tested in both assays (Figure 8B and C) for antiviral activity and their cytotoxicity was also evaluated. While destruxin A and B resulted as non-active against HCoV-OC43 infection in MRC-5 cells and showed cytotoxicity in the range of 10 µM, linoleic acid and gymnoascolide A exhibited EC_50_ values of 18.31 and 20.62 μM, respectively, with no evident toxicity against MRC-5 host cells (Figure 8B and C). Additionally, we evaluated by RT-PCR the presence of HCoV-OC43 in culture supernatants after 72h of incubation with gymnoascolide A and linoleic acid (Figure 8D). Gymnoascolide A at 80 µM reduced the viral load in the supernatant below 5% and at 40 μM HCoV-OC43 RNA levels remained under 20%. For linoleic acid, the HCoV-OC43 RNA load was below 50% at a 15 μM concentration.

**Figure 8.**
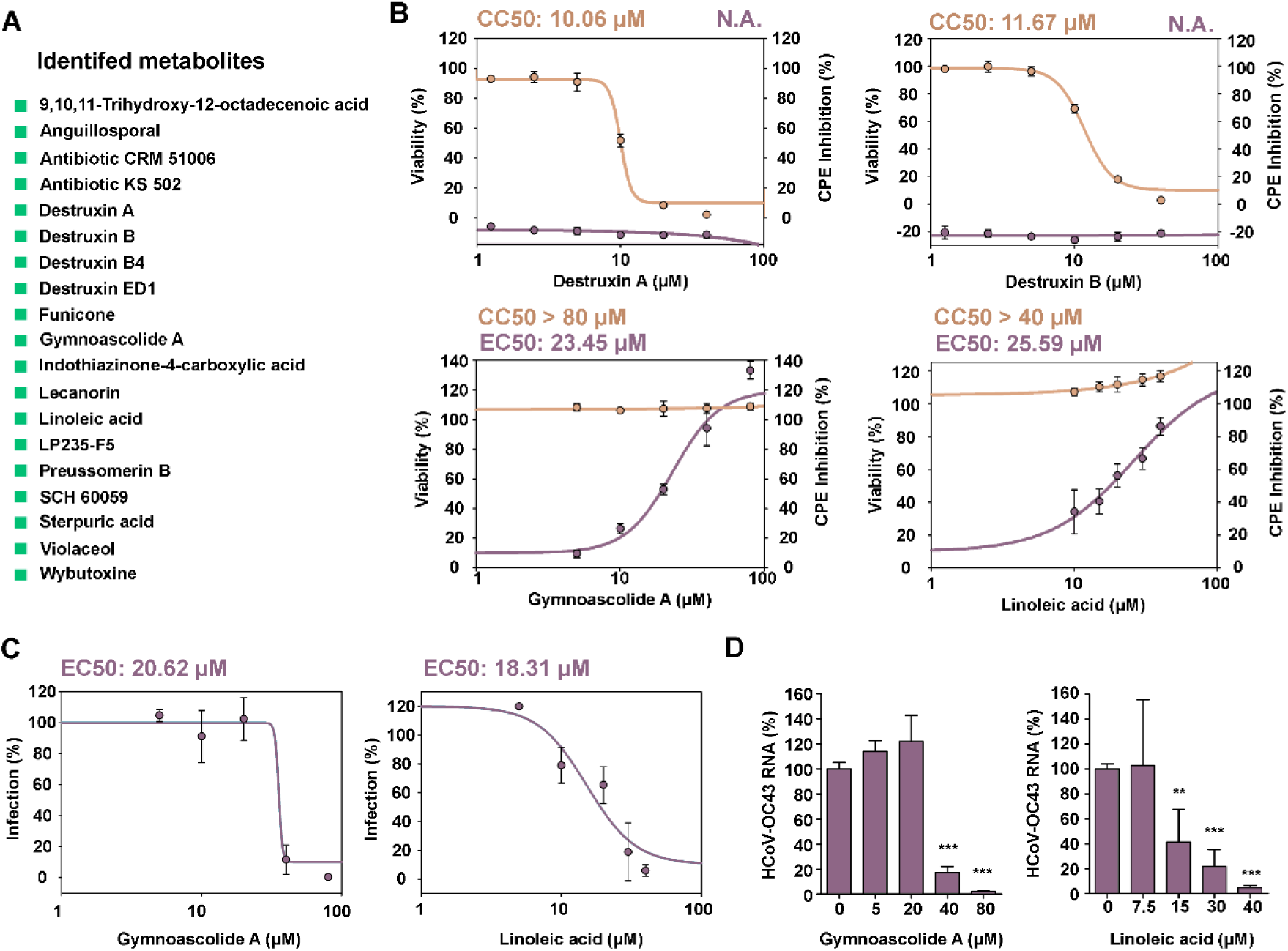
Antiviral activity and cytotoxicity evaluation of the known metabolites identified by LC-MS. A. List of the 19 identified known compounds in LC-MS analysis. B. Dose-response curve for CPE inhibition at 5 days post-infection and 96 h in the presence of each drug. EC_50_ values for the CPE inhibition assay and CC_50_ values for MRC-5 cells. C. Dose response curves obtained with the HCS assay at 4 days post-infection and 72 h in the presence of the drug. EC_50_ value for the HCS assay. D. Extracellular HCoV-OC43 RNA levels after 72h exposure to gymnoascolide A and linoleic acid. EC_50_ and CC_50_ values were calculated using SigmaPlot software.

After LC-MS dereplication, 15 extracts containing potentially novel components and showing well-resolved metabolic profiles were prioritized for small scale regrowth. The antiviral activity was confirmed for 10 of them.

## DISCUSSION

The outbreaks of coronavirus infections that cause high case-fatality rates in the last decade since the SARS-CoV-1 outbreak in 2003 followed by MERS in 2012 and SARS-CoV-2 in 2020 have highlighted the inadequacy of existing antiviral treatments. Together with vaccines, drugs are essential to battle future epidemics making necessary the discovery of new antivirals. Microbial natural products are an interesting alternative for drug discovery due to their chemical diversity. The main purpose of this work was to establish a robust HTS platform for the discovery of new antivirals from microbial extracts that may serve as broad-spectrum antivirals for future outbreaks. Fundación MEDINA has available for screening the world’s largest collection of microbial natural product extracts harboring over 200,000 microbial natural products extracts generated from over 190,000 strains of filamentous fungi, actinomycetes and unicellular bacteria cultured in diverse formats to ensure a rich metabolite-producing spectrum. This collection has been thoroughly annotated with both chemical and biological data for tailor-made drug discovery purposes. In the current proof-of-concept (pilot) campaign, we screened 1280 extracts, 390 of them from fungi and 960 of from actinomycetes. The use of natural products collections requires the expertise to isolate and identify the pure compound responsible for the antiviral activity. The workflow (Figure 3) used in this platform guarantees that only those extracts tested and rigorously proven to contain activity and potentially new metabolites are progressed.

We have successfully implemented two different methodologies to assess antiviral activity of microbial crude extracts against HCoV-OC43. The host selected for HCoV-OC43 propagation is a human lung fibroblast cell line, MRC-5. HCoV-OC43 infection in MRC-5 cells produced a more pronounced cytopathic effect after 5 days of infection when compared with hamster cells BHK-21. In a recent work Schirtzinger et al, also found MRC-5 to be the most suitable host for HCoV-OC43 infection when compared to other cells lines commonly used as viral hosts (13). The plaque assay, widely used in viral studies, was not suitable for HCoV-OC43 infection of MRC-5; despite HCoV-OC43 being a lytic virus, the cell morphology made the plaques grow into each other (13). In addition to the plaque and antibody-based assays for TCID50 determination, resazurin reduction is a fluorimetric assay that has also been used in antiviral and neutralization assays when evaluating lytic virus infection (30, 31).

Fluorimetric-based assays have some limitations. In the case of resazurin, since its conversion depends upon an enzymatic reaction, certain compounds may interfere by directly inhibiting this process. Likewise, coloured compounds may interfere with the assay (30). To overcome the possible limitations, we set up an HCS assay that allows us to simultaneously evaluate both antiviral activity and possible toxicity of different sets of compounds by quantifying indirect immunostaining of the HCoV-OC43 nucleocapsid and MRC-5 cells nuclei staining.

We therefore have established a new and standardised methodology for the evaluation and quantification of HCoV-OC43 infection upon treatment with either pure compounds or microbial natural extracts that may have potential antiviral activity. The methodology was first validated with three compounds that had already been described as antivirals: chloroquine, mycophenolate mofetil and ribavirin (25-28). All three compounds showed activity against HCoV-OC43 in both assays. Although mycophenolate mofetil showed promising results, in a previous study severe side effects in an *in vivo* model in marmosets infected with MERS (32) and no antiviral activity against HCoV-OC43 in a mouse model were described (26).

Our efficient LC-MS dereplication methodology allows us to single out active extracts containing known metabolites. The identification of novel compounds is key to develop new antiviral treatments; however, we should not overlook known compounds that could provide a starting point in the development of new therapies. Destruxins A and B, were previously described antivirals against hepatitis B virus (HBV), although they did not have an effect on HCoV-OC43 infection (7, 33). On the other hand, linoleic acid and gymnoascolide A showed activity against HCoV-OC43. While this is the first time gymnoascolide A is described to have anticoronavirus activity, different *in silico* studies have related linoleic acid to interact and interfere with SARS-CoV-2. Toeltzer et al. were the first to describe the interaction of linoleic acid with the RBD of the spike protein of SARS-CoV-2 (34). In a recent study, linoleic acid was reported to interact and inhibit RNA dependent RNA polymerase (RdRp) *in silico*. They also performed *in vitro* and *in vivo* studies, where they exposed the cells to linoleic acid at different time points of the infection. In both models they observed an antiviral effect of linoleic acid with the reduction of viral RNA load in the media and viral RNA present in tissues respectively (35).

The confirmation of the antiviral activity of linoleic acid and gymnoascolide A validates the strategy used in this screening approach. From the LC-MS dereplication data we can conclude that our screening platform is able to identify known compounds from crude extracts with antiviral activity, proving that we can dereplicate extracts containing cytotoxic metabolites or known compounds that have not been previously described as antivirals. The drawback of this methodology is that we may miss active metabolites that are less abundant than the cytotoxic ones.

This is the first report of an established HCS methodology for the discovery of new anti-HCoV-OC43 antivirals present in complex microbial extracts. From a collection of 1280 extracts we have identified and confirmed 10 different fungal extracts with new potent anti-HCoV-OC43 activity. These hits are currently undergoing further studies to identify the metabolites responsible for the antiviral activity. We have shown that this platform can be used to validate already pure compounds with previously described antiviral activity such as ribavirin or linoleic acid, but also for the discovery of novel antiviral activity of known natural products, as in the case of gymnoascolide A.

## SUPPLEMENTAL FIGURES

**Supplemental Figure 1.**
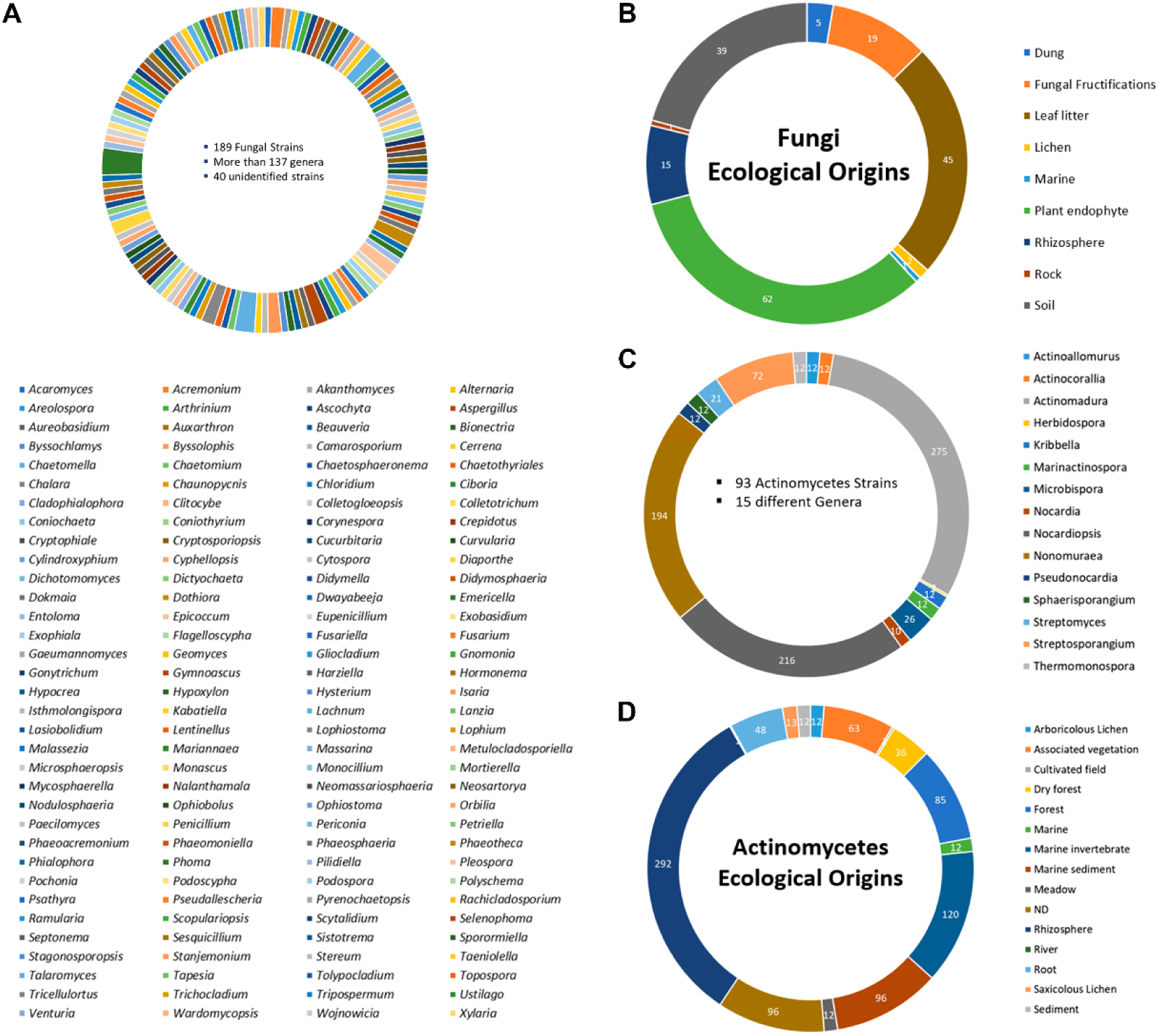
Selected 1280 microbial extracts from MEDINA’s Natural Product library. A. Genera of fungal extracts. B. Ecological origins of the 189 fungal strains. C. Genera of actinomycetes extracts. D. Ecological origin of the 93 actinomycetes strains.

## ACKNOWLEDGEMENTS

This work was funded by the the European Commission—Next Generation EU (regulation EU 2020/2094), through CSIC’s Global Health Platform (PTI Salud Global), the Instituto de Salud Carlos III Subdirección General de Redes y Centros de Investigación Cooperativa-Red de Investigación Cooperativa en Enfermedades Tropicales (RICET: RD16/0027/0014), the MCIN/AEI/10.13039/501100011033 (PID2019-109623RB-I00), the MCIN/AEI/10.13039/501100011033 and FEDER Una manera de hacer Europa (2016-79957-R) and by the Junta de Andalucía (BIO-199).

